# Septins regulate heart contractility through modulation of cardiomyocyte store-operated calcium entry

**DOI:** 10.1101/2024.11.04.621876

**Authors:** Benjamin A. Tripoli, Jeremy T. Smyth

## Abstract

Highly regulated cardiomyocyte Ca^2+^ fluxes drive heart contractions. Recent findings from multiple organisms demonstrate that the specific Ca^2+^ transport mechanism known as store-operated Ca^2+^ entry (SOCE) is essential in cardiomyocytes for proper heart function, and SOCE dysregulation results in cardiomyopathy. Mechanisms that regulate SOCE in cardiomyocytes are poorly understood. Here we tested the role of cytoskeletal septin proteins in cardiomyocyte SOCE regulation. Septins are essential SOCE modulators in other cell types, but septin functions in cardiomyocytes are nearly completely unexplored. We show using targeted genetics and intravital imaging of heart contractility in *Drosophila* that cardiomyocyte-specific depletion of septins 1, 2, and 4 results in heart dilation that phenocopies the effects of SOCE suppression. Heart dilation caused by septin 2 depletion was suppressed by SOCE upregulation, supporting the hypothesis that septin 2 is required in cardiomyocytes for sufficient SOCE function. A major function of SOCE is to support SERCA-dependent sarco/endoplasmic reticulum (S/ER) Ca^2+^ stores, and augmenting S/ER store filling by SERCA overexpression also suppressed the septin 2 phenotype. We also ruled out several potential SOCE-independent septin functions, as septin 2 phenotypes were not due to septin function during development and septin 2 was not required for z-disk organization as defined by α-actinin labeling. These results demonstrate, for the first time, an essential role of septins in cardiomyocyte physiology and heart function that is due, at least in part, to septin regulation of SOCE function.

## INTRODUCTION

Contractions of cardiomyocytes, the muscle cells of the heart, are driven by cytoplasmic Ca^2+^ transients that are generated through a process known as excitation-contraction coupling (ECC). During excitation-contraction coupling, depolarization of the plasma membrane or sarcolemma activates voltage-gated L-type Ca^2+^ channels, allowing a small quantity of Ca^2+^ to enter the cytoplasm. This Ca^2+^ from L-type channels then directly activates ryanodine receptor (RyR) Ca^2+^ release channels in the sarcoplasmic reticulum (SR), resulting in a large cytoplasmic Ca^2+^ signal due to Ca^2+^ efflux from SR Ca^2+^ stores. It is the large quantity of Ca^2+^ that is released from the SR that drives myofilament sliding and cardiomyocyte contraction (Bers, 2002). Proper regulation of SR Ca^2+^ stores is therefore essential to physiological heart contractility, and several cardiomyopathies and heart failure are strongly associated with SR Ca^2+^ store dysregulation (Adeniran et al., 2015; Bers, 2002; Eisner et al., 2020; Njegic et al., 2020). Stored Ca^2+^ that is released by the RyR with each ECC cycle is pumped back into the SR by sarco/endoplasmic reticulum ATPase (SERCA) pumps (Kho, 2023). However, any net loss of Ca^2+^ from the cell can limit the available Ca^2+^ for SERCA pumps and result in SR Ca^2+^ store depletion. Store-operated Ca^2+^ entry (SOCE) is a Ca^2+^ influx mechanism that is activated specifically in response to SR Ca^2+^ store depletion, and Ca^2+^ that enters the cell via SOCE can be pumped into the SR by SERCA to replenish SR Ca^2+^ stores (Gruszczynska-Biegala et al., 2011; Ong et al., 2019). Recent work from our lab and others has demonstrated an essential role for cardiomyocyte SOCE in normal heart physiology, as cardiomyocyte SOCE suppression results in dilated cardiomyopathy and compromised cardiac output (Collins et al., 2014; Parks et al., 2016; Petersen et al., 2020). However, the mechanisms that regulate SOCE in cardiomyocytes and the specific functions of SOCE that are required for normal heart physiology are still poorly understood.

SOCE is a near-ubiquitous process that is functional in most animal cell types. SOCE is best understood in non-excitable cells such as immune and secretory cells, where it is coupled to endoplasmic reticulum (ER) Ca^2+^ store release by inositol 1,4,5-trisphosphate receptors (IP_3_R) (Emrich et al., 2022). The role of SOCE in excitable cells including muscle is less clear. The SOCE Ca^2+^ influx mechanism consists of two primary molecular components: stromal interaction molecules, or Stim proteins, which function as SR Ca^2+^ sensors, and Orai Ca^2+^ influx channels in the plasma membrane. Stim proteins are located in the E/SR membrane and have an N-terminal EF-hand Ca^2+^ binding domain within the E/SR lumen. SR Ca^2+^ store depletion results in Ca^2+^ dissociation from Stim’s EF-hand domain, causing a conformational change that allows Stim to directly interact with and activate Orai Ca^2+^ influx channels. This interaction between Stim and Orai occurs at contact sites between the E/SR and plasma membranes (Putney, 2018). Mammals express two Stim molecules, STIM1 and STIM2, and three Orais (Orai1, Orai2, and Orai3), with STIM1 and Orai1 being the most widely expressed isoforms in most tissues. An accumulation of studies from diverse species now clearly demonstrate that SOCE plays a vital role in supporting normal heart function. For example, cardiomyocyte-specific deletion of STIM1 results in left ventricular dilation and reduced cardiac output in mice (Collins et al., 2014; Parks et al., 2016). Orai3 deletion also results in dilated cardiomyopathy and heart failure in mice (Gammons et al., 2021), while Orai1 suppression results in reduced cardiomyocyte fractional shortening and heart failure in zebrafish (Volkers et al., 2012). Our own work in *Drosophila* also demonstrates that cardiomyocyte-specific depletion of the single Stim or Orai isoforms expressed in these animals causes dilated cardiomyopathy and early lethality (Petersen et al., 2020). A key question that remains to be addressed is how is SOCE properly regulated in cardiomyocytes to support physiological heart contractility and function? Mounting evidence from other cell types demonstrates that members of the septin family of cytoskeletal GTPases are essential SOCE regulators (Deb and Hasan, 2016), but the functional role of septins in cardiomyocytes is almost completely unknown.

Mammalian septins are subdivided into four groups of functionally homologous members: SEPT2 (SEPT1, SEPT2, SEPT4, SEPT5), SEPT3 (SEPT3, SEPT9, SEPT12), SEPT6 (SEPT6, SEPT8, SEPT10, SEPT11, SEPT14), and SEPT7 (SEPT7). Mammalian septins hetero-oligomerize to form hexameric or octameric filaments with the subunit group order SEPT2-SEPT6-SEPT7-SEPT7-SEPT6-SEPT2 for hexamers and SEPT2-SEPT6-SEPT7-SEPT3-SEPT3-SEPT7-SEPT6-SEPT2 for octamers (Cavini et al., 2021). Septin filaments generally localize to membranes and they are essential for numerous cellular processes including cytokinesis, ciliogenesis, and phagocytosis (Woods and Gladfelter, 2021). A role for septins in SOCE regulation was first demonstrated through a genome-wide siRNA screen in HeLa cells, in which it was shown that depletion of SEPT2 group members SEPT4 and SEPT5 results in suppression of SOCE-mediated Ca^2+^ influx (Sharma et al., 2013). Subsequent studies have demonstrated that septins modulate SOCE function by regulating ER-plasma membrane contact sites and Stim-Orai interaction at these sites (Katz et al., 2019; Sharma et al., 2013), as well as by regulating specific actin organizations at the plasma membrane that are required for Stim clustering and Orai activation (de Souza et al., 2021). The requirement for septins for proper SOCE activation has also been supported by a number of studies in *Drosophila*. *Drosophila* express five highly conserved septin subunits, with *Drosophila* Sept1 and Sept4 orthologous to mammalian SEPT2 group members, *Drosophila* Sept2 and Sept5 orthologous to SEPT6 group members, and *Drosophila* Sept7 or Pnut orthologous to SEPT7 (Shuman and Momany, 2021). The lack of SEPT3 orthologs means that *Drosophila* septins only form hexamers. Similar to mammalian cells, depletion of *Drosophila* SEPT2 subunits Sept1 and Sept4 strongly attenuated SOCE in neurons (Deb et al., 2016). This finding in *Drosophila* was also extended to SEPT6 subunits, as *Drosophila* Sept2 depletion also resulted in neuronal SOCE attenuation (Deb and Hasan, 2019). Thus, septin subunits from the SEPT2 and SEPT6 groups have essential and highly conserved roles in positive SOCE regulation in both excitable and non-excitable cells.

The goals of this project were to test the role of cardiomyocyte septins in heart function and to determine if there is functional association of septins with the SOCE pathway in these cells. We carried this out using genetic tools in the *Drosophila* heart. The *Drosophila* heart is a tube-shaped organ composed of contractile cardiomyocytes, and it circulates a lymph-like fluid called hemolymph throughout the animal’s body in an open circulatory system (Souidi and Jagla, 2021). Importantly, contractile physiology, including excitation-contraction coupling, is highly conserved between *Drosophila* and mammalian cardiomyocytes (Lin et al., 2011).

Cardiomyocyte SOCE suppression in *Drosophila* results in phenotypes similar to those in mammals (Petersen et al., 2020), suggesting that SOCE function in *Drosophila* cardiomyocytes is also well-conserved. Thus, the *Drosophila* heart is a powerful model for testing conserved septin functions in cardiomyocyte SOCE regulation. Our results demonstrate that cardiomyocyte depletion of SEPT2 and SEPT6 group subunits results in dilated cardiomyopathy that phenocopies cardiomyocyte Stim and Orai depletion. We further demonstrate that the septin phenotypes are suppressed by genetic upregulation of SOCE function as well as by SERCA overexpression. These results demonstrate, for the first time, a functional role for SEPT2 and SEPT6 subunits in cardiomyocyte physiology that is due, at least in part, to positive regulation of SOCE.

## RESULTS

### Silencing Septins 1, 2, or 4 in cardiomyocytes causes cardiac dilation similar to SOCE suppression

We first tested the effect of septin 1, 2, and 4 suppression on heart function in 3-5 day old adult flies using cardiomyocyte-specific RNAi expression driven by *tinC-GAL4* (Ocorr et al., 2014). We used a cardiomyocyte-specific RNAi approach due to availability of reagents and to avoid cardiac-independent effects of whole-animal mutants. These animals also had cardiomyocyte-restricted tdTomato (CM-tdTom) expression for intravital imaging of heart contractility (Klassen et al., 2017; Petersen et al., 2020). We found that individual suppression of septins 1, 2, and 4 all resulted in significant heart dilation, as can be seen in representative still-frame images of diastole and systole and representative M-mode traces (Figure 1A, B). Direct measurements of end-diastolic dimensions (EDD) and end-systolic dimensions (ESD) from the M-mode traces demonstrated significant increases in both EDD and ESD for each of the septin groups, confirming marked dilation caused by septin suppression (Figure 1C, D). Fractional shortening (FS), a quantitative indicator of systolic function and cardiac output, was also significantly decreased by approximately 15-20% in hearts with cardiomyocyte-specific septin suppression (Figure 1E). We observed the significant changes in EDD, ESD, and FS in the septin-suppressed versus control hearts in both male and female animals, though the effects were somewhat more pronounced in females (Supplementary Figure S1). We therefore focused on female animals for the remainder of our studies. Heart rate and rhythmicity were not significantly altered in the septin-suppressed hearts compared to controls (Supplementary Figure S2). We also replicated our results with septin 1, 2, and 4 suppressions by RNAi using a second driver that expresses in cardiomyocytes, *hand4.2-GAL4* (Han et al., 2006), suggesting the phenotypes we observed are due to cardiomyocyte-specific septin suppression (Supplementary Figure S3). We next took advantage of an available Drosophila septin 2 loss-of function mutant, *septin2^2^*. As shown previously, homozygous *septin2^2^* animals exhibit significant developmental lethality but it is possible to obtain adult escapers, allowing us to analyze adult heart function in these animals (O’Neill and Clark, 2013; O’Neill and Clark, 2016). Consistent with the RNAi results, EDD and ESD were significantly increased and FS significantly decreased in *septin2^2^*compared to *septin2^2^*/*+* or *w^1118^* control hearts (Figure F,-I). This important result confirms the essential requirement for septin 2 for proper heart function and suggests that the RNAi results are specific to septin suppression and not due to off-target effects. Overall, the combination of increased EDD and ESD and decreased FS in septin-suppressed hearts is consistent with the clinical features of dilated cardiomyopathy. And importantly, these results closely phenocopied the effects of cardiomyocyte-specific *Stim* suppression, which also resulted in increased EDD and ESD and decreased FS (Figure J-M). This is consistent with the hypothesis that the effects of septin suppression on the heart are due to suppression of Stim-dependent SOCE function.

**Figure 1.**
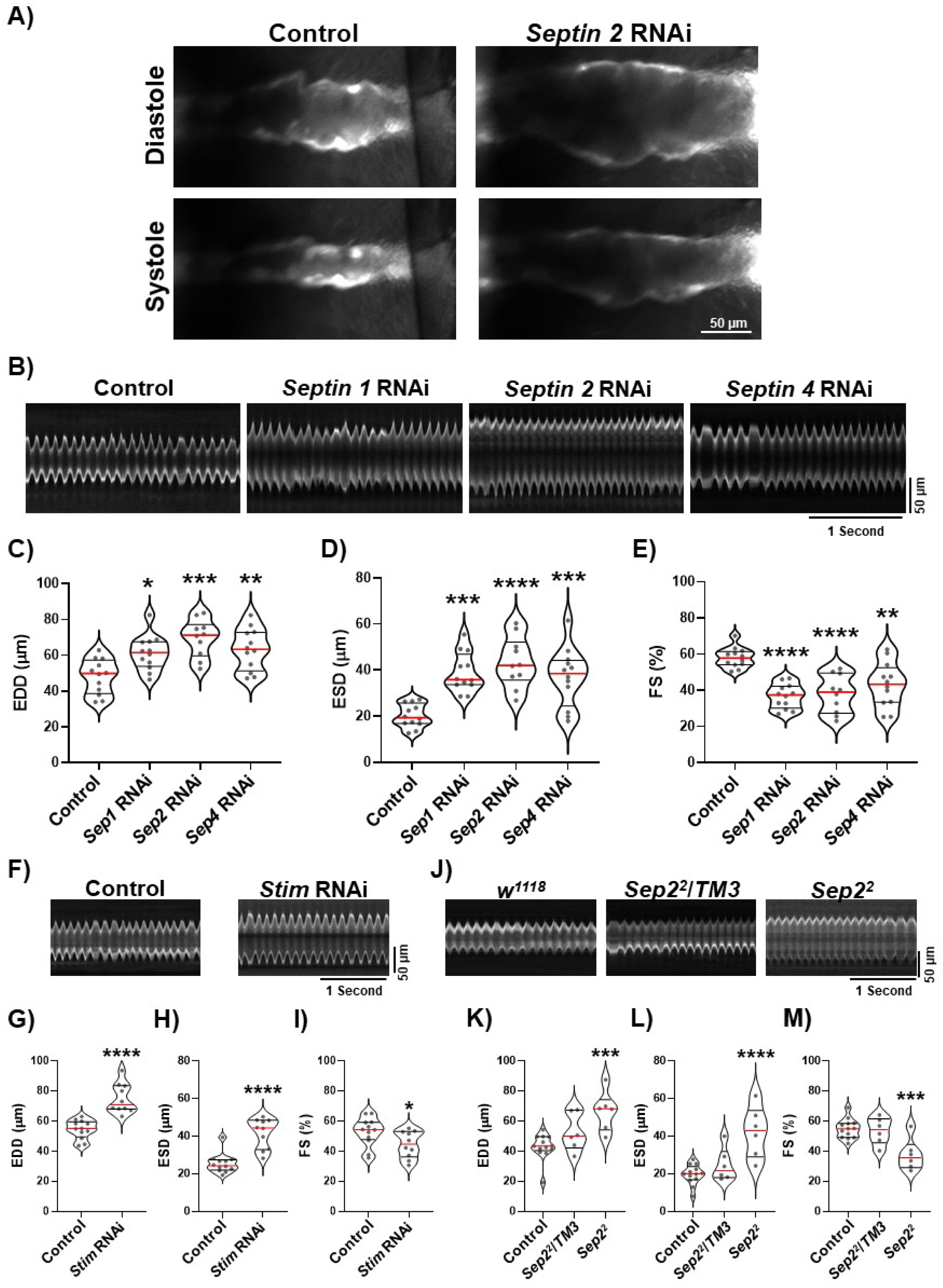
Septin 1, 2, and 4 suppression results in heart dilation. A) Representative still-frame images of CM-tdTom expressing *Drosophila* hearts at diastole and systole with *tinC-GAL4* driven control or *septin 2* RNAi. Images were taken from 5 second timelapse image sequences acquired at 200 frames/second. Note the significant increase in width of the *septin 2* RNAi compared to control hearts at both diastole and systole. B) Representative M-mode traces from animals with *tinC-GAL4* driven control, *septin 1*, *septin 2*, or *septin 4* RNAi as indicated. C-E) Plots of EDD, ESD, and FS, respectively, calculated from M-modes for animals with indicated *tinC-GAL4* driven RNAi. F) Representative M-modes from animals with *tinC-GAL4* driven control or *Stim* RNAi. G-I) Plots of EDD, ESD, and FS, respectively, calculated from M-modes for animals with indicated *tinC-GAL4* driven RNAi. J) Representative M-modes from control (*w^1118^*), *Sep2^2^* heterozygous (*Sep2^2^/TM3*), or *Sep2^2^* homozygous animals. K-M) Plots of EDD, ESD, and FS, respectively, calculated from M-modes for animals with indicated genotypes. For all plots each symbol represents the average of five measurements from a single animal’s M-mode. Red lines indicate median and black lines indicate quartiles. *, P<0.05; **, P<0.01, ***, P < 0.001; ****, P < 0.0001 compared to control; one-way ANOVA with Tukey’s multiple comparisons for C-D and K-M, two-tailed t-test for G-I.

### SOCE activation suppresses dilation of septin 2-deficient hearts

Given the striking similarities of the Septin and Stim suppression phenotypes and the previously defined role for septins in SOCE regulation, we hypothesized that septin suppression may cause heart dilation due to insufficient SOCE activation. We tested this by driving constitutive SOCE in septin 2-depleted hearts using constitutively active *Orai* and *Stim* mutant transgenes. We focused on septin 2 for the remainder of our experiments because of the strong phenotypes with septin 2 suppression and the lack of redundancy with other septin subunits that complicates analysis of septins 1 and 4. Constitutively active Orai has a glycine to methionine substitution in the channel hinge that locks the channel in an open conformation (Zheng et al., 2013). We previously demonstrated that this gain-of-function *Orai* mutant causes hypertrophy of the *Drosophila* heart (Petersen et al., 2022), consistent with cardiac hypertrophy that results from gain of SOCE function in mammals (Hulot et al., 2011). We found that the enlarged systolic dimensions of *septin 2* RNAi hearts were reversed to near-control values when *Orai^CA^*was co-expressed, and reduced FS of *septin 2* suppressed hearts was also reversed (Figure 2A-D). Enlarged diastolic dimensions of *septin 2* suppressed hearts, on the other hand, were not as markedly reversed by *Orai^CA^*expression.

**Figure 2.**
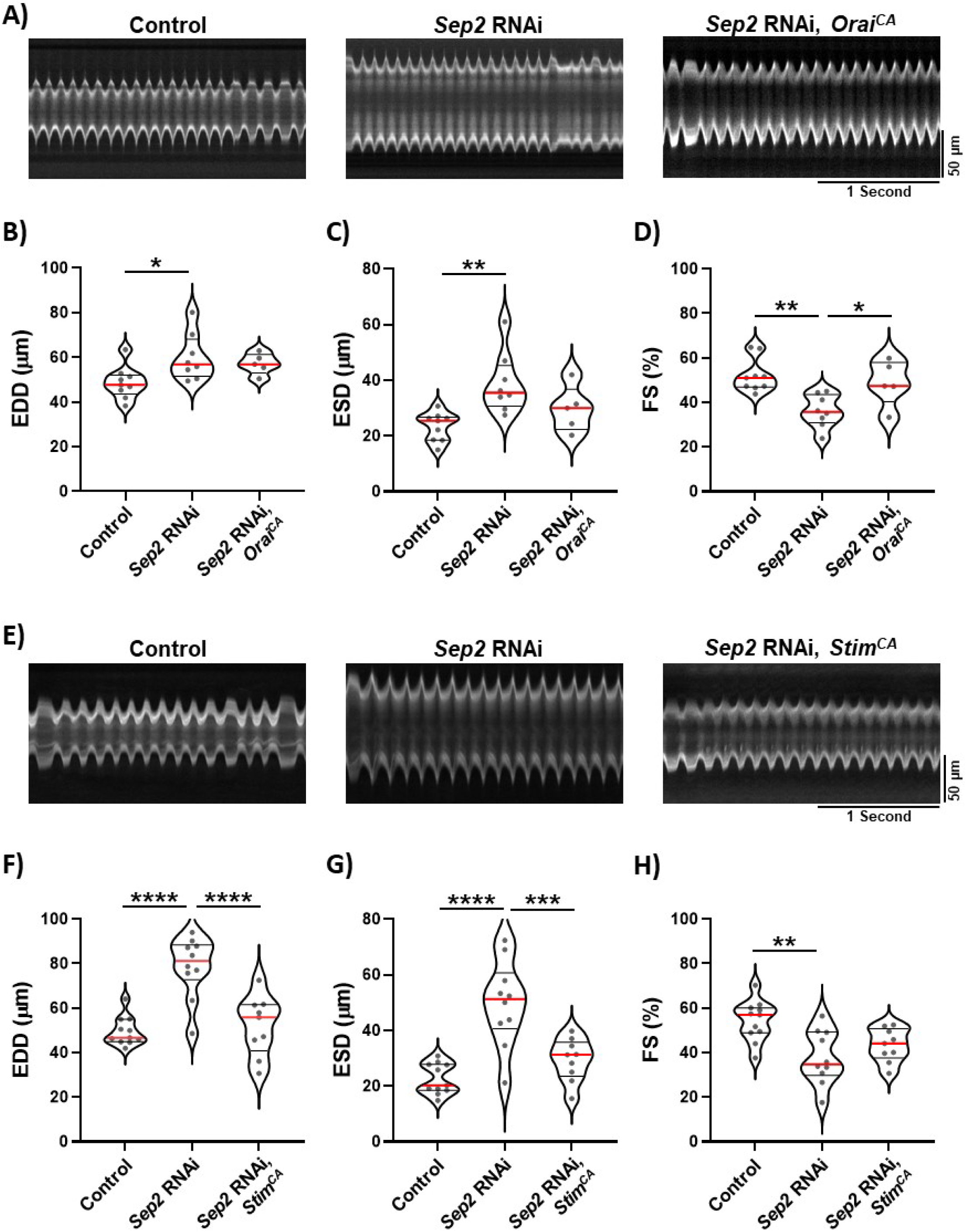
Septin 2 phenotypes are suppressed by SOCE upregulation. A) Representative M-mode traces from animals with *tinC-GAL4* driven control RNAi, *septin 2* RNAi, or *septin 2* RNAi with Orai^CA^. B-D) Plots of EDD, ESD, and FS, respectively, calculated from M-modes for animals with indicated *tinC-GAL4* driven transgenes. E) Representative M-mode traces from animals with *tinC-GAL4* driven control RNAi, *septin 2* RNAi, or *septin 2* RNAi with Stim^CA^. F-H) Plots of EDD, ESD, and FS, respectively, calculated from M-modes for animals with indicated *tinC-GAL4* driven transgenes. Each symbol in plots represents the average of five measurements from a single animal’s M-mode. Red lines indicate median and black lines indicate quartiles. *, P<0.05; **, P<0.01, ***, P < 0.001; ****, P < 0.0001 compared to control; one-way ANOVA with Tukey’s multiple comparisons.

We next tested constitutively active Stim, which has two aspartic acid to alanine substitutions in the Ca^2+^-sensing EF-hand that lock Stim in a Ca^2+^-depleted, active conformation (Zhang et al., 2005). This *Stim^CA^* transgene also drives hypertrophy of the *Drosophila* heart (Petersen et al., 2022). We found that *Stim^CA^* expression strongly reversed enlargement of both diastolic and systolic dimensions caused by *septin 2* depletion, and also restored FS to near control vales (Figure 2E-H). These results demonstrate that gain of SOCE function can largely reverse the effects of *septin 2* depletion, supporting the hypothesis that the septin 2 effects are at least partly due to SOCE suppression. Differences in the reversal of septin 2 phenotypes with Orai^CA^ versus Stim^CA^ may be due to stronger expression and/or penetrance of the Stim^CA^ transgene.

### SERCA overexpression suppresses dilation of STIM, septin 2 deficient hearts

A major function of SOCE is to provide cytoplasmic Ca^2+^ to SERCA pumps to refill SR Ca^2+^ stores. Importantly, dysregulation of cardiomyocyte SR Ca^2+^ uptake is strongly associated with heart failure (Luo and Anderson, 2013). It is therefore possible that the cardiomyocyte septin 2 depletion phenotypes may ultimately be caused by a reduction in SR Ca^2+^ store content due to suppression of SOCE. We tested this by determining whether SERCA overexpression, which would be expected to increase SR Ca^2+^ content, can reverse the septin 2 depletion phenotypes. It was first important to determine whether SERCA overexpression can reverse the effects of SOCE suppression. In support of this, we found that the significant increases in EDD and ESD caused by cardiomyocyte Stim depletion were fully reversed by SERCA overexpression (Figure 3A-C). Decreased FS with Stim depletion was also strongly reversed by SERCA overexpression (Figure 3D). SERCA overexpression alone did not significantly affect EDD, ESD, or FS compared to controls (Figure 3B-D). Thus, cardiomyocyte SERCA overexpression is an effective strategy for reversing the deleterious effects of SOCE suppression on heart function. We then found that similar to the results for Stim, increased EDD and ESD and decreased FS caused by septin 2 depletion were also fully reversed by SERCA overexpression (Figure 3A-D). These results strongly support our hypothesis that septin 2 is required in cardiomyocytes to support SOCE function that is essential for maintaining SR Ca^2+^ store content and proper heart contractility. We attempted to further support these findings by directly measuring both SR Ca^2+^ store content and contractile Ca^2+^ transients in *Drosophila* cardiomyocytes based on imaging of the genetically encoded jRGECO Ca^2+^ indicator *in vivo* and in partially dissected preparations. Unfortunately, however, technical challenges prevented us from acquiring reliable data from these experiments.

**Figure 3.**
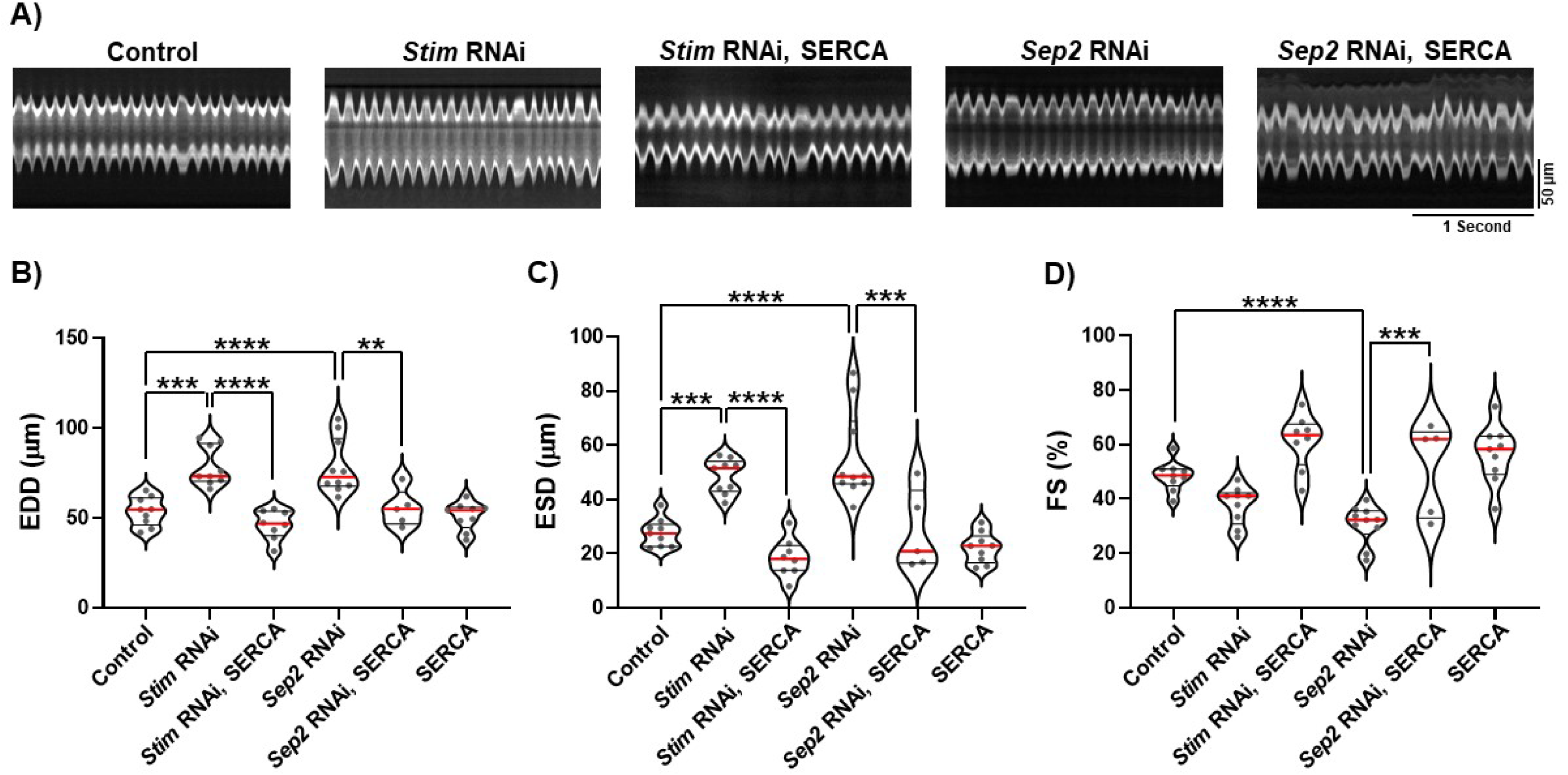
Septin 2 phenotypes are suppressed by SERCA overexpression. A) Representative M-mode traces from animals with *tinC-GAL4* driven transgenes as indicated. B-D) Plots of EDD, ESD, and FS, respectively, calculated from M-modes for animals with indicated *tinC-GAL4* driven transgenes. Each symbol in plots represents the average of five measurements from a single animal’s M-mode. Red lines indicate median and black lines indicate quartiles. **, P<0.01, ***, P < 0.001; ****, P < 0.0001; one-way ANOVA with Tukey’s multiple comparisons.

### Septin 2 phenotypes do not involve developmental defects or alterations of z-disc integrity

Our results thus far suggest that a major function of septin 2 in cardiomyocytes is SOCE regulation. We also considered the possibility that septin 2 has additional, SOCE-independent functions in cardiomyocytes that are required for proper heart physiology. Septins are required for cytokinesis and because cardiomyocytes in the adult heart are post-mitotic, cytokinesis dysfunction due to septin 2 depletion in cardiomyocytes would most likely occur during heart development. We therefore tested whether septin 2 depletion phenotypes in the adult heart are due to developmental dysfunctions by initiating cardiomyocyte septin 2 depletion post-developmentally in adults using the *GAL4*/*GAL80[ts]* inducible expression system. *GAL80* is a competitive *GAL4* inhibitor that blocks transcriptional activation when active. Temperature sensitive *GAL80* (*GAL80[ts]*) is active at the permissive temperature of 18° C but is inactive at temperatures above 27° C. Thus, animals co-expressing *GAL80[ts]* and *GAL4* are reared at 18° C to prevent *UAS* gene expression by *GAL4*, but moved to 27° C to induce expression. We generated animals that co-express ubiquitous *tubP-GAL80[ts]* with *tinC-GAL4*, together with either septin 2 or control RNAi. These animals also expressed CM-tdTom for heart contractility analysis. We first tested animals that were maintained at 18° C throughout development as well as adulthood until the time of analysis as a control to verify *GAL80[ts]* efficacy, as this temperature paradigm should suppress RNAi expression for the entire experiment. As expected, EDD, ESD, and FS values were no different for septin 2 versus control RNAi animals maintained at 18° C throughout the experiment (Figure 4A-D). We then tested animals that were maintained at 18° C throughout development but were then shifted to 27° C as one or two day-old adults to initiate RNAi expression. Heart contractility was then analyzed in these animals 3-5 days after the shift to 27° C. Remarkably, EDD and ESD values of temperature-shifted *septin 2* RNAi animals were significantly increased compared to control RNAi animals (Figure 4E-G), similar to results in Figure 1 with RNAi expression throughout development. FS was slightly decreased in *septin 2* versus control RNAi animals but this was not statistically significant (Figure 4H). These results essentially recapitulate the effects of septin 2 depletion throughout development, suggesting that any cardiomyocyte septin 2 functions during development do not contribute significantly to the contractility phenotypes we observe in the adult heart.

**Figure 4.**
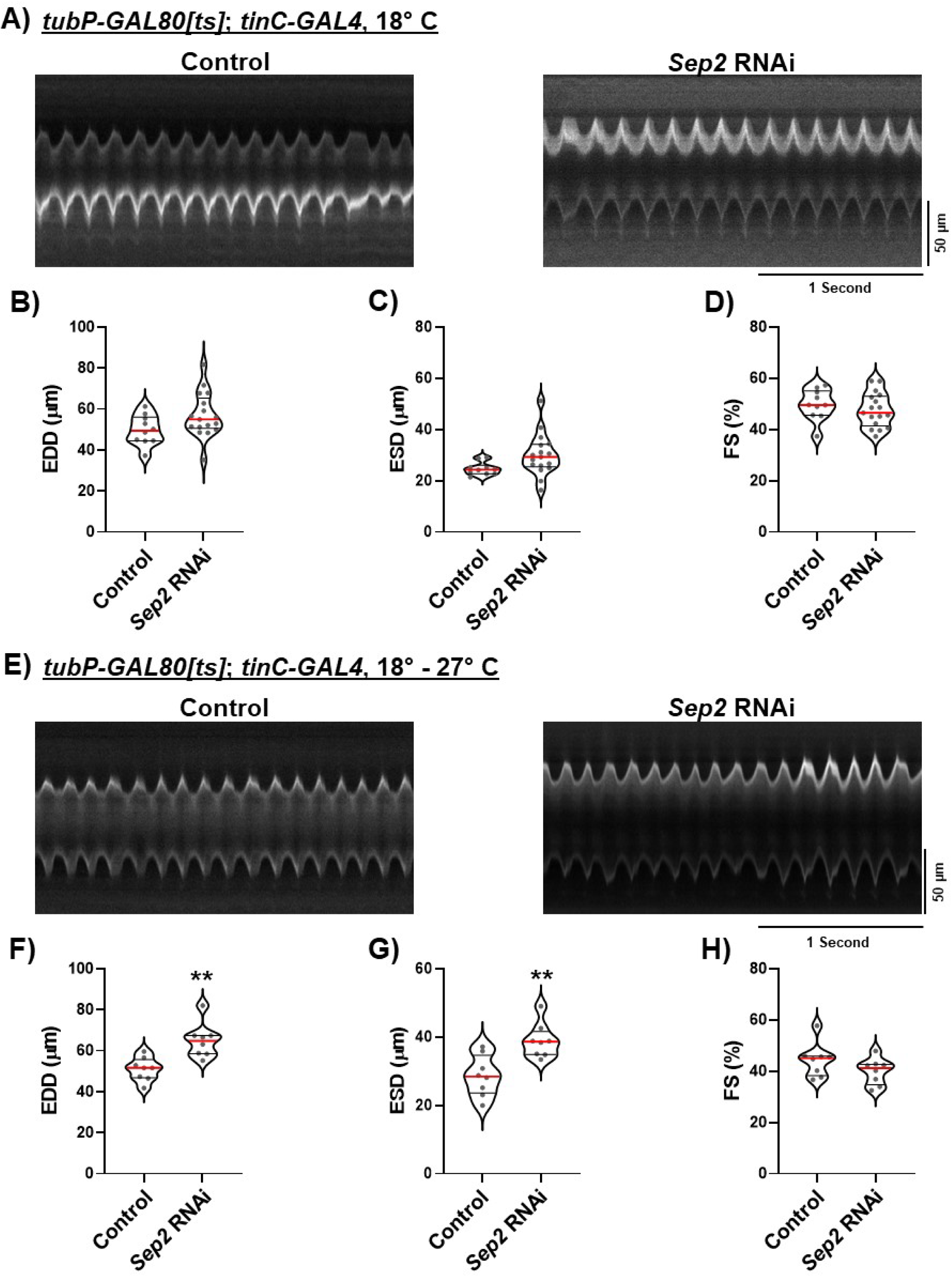
Post-developmental septin 2 depletion is sufficient to drive the heart dilation phenotype. A) Representative M-mode traces for animals expressing *TubP-GAL8[ts]* and *tinC-GAL4* with either control or *septin 2* RNAi. These animals were maintained at 18° C to suppress RNAi expression from the time of egg laying until intravital imaging was carried out as adults. B-D) Plots of EDD, ESD, and FS, respectively, calculated from M-modes for animals expressing *TubP-GAL80[ts]* and *tinC-GAL4* and indicated RNAi and maintained continuously at 18° C until intravital imaging as adults as described in A. E) Representative M-mode traces for animals expressing *TubP-GAL80[ts]* and *tinC-GAL4* with either control or *septin 2* RNAi. These animals were first maintained at 18° C from the time of egg laying until adults were one day old to suppress RNAi expression throughout development. One day adults were then moved to 27° C to activate RNAi expression, and were maintained at 27° C for 3-5 days until intravital imaging was carried out. F-H) Plots of EDD, ESD, and FS, respectively, calculated from M-modes for animals expressing *TubP-GAL80[ts]* and *tinC-GAL4* and indicated RNAi and moved from 18° C to 27° C as one day old adults as described in E. Each symbol in plots represents the average of five measurements from a single animal’s M-mode. Red lines indicate median and black lines indicate quartiles. **, P<0.01 compared to control, two-tailed t-test.

It has previously been shown that septin 7 localizes to z-disks in mouse skeletal myofibers (Gönczi et al., 2022), and that septin 7 depletion in zebrafish results in z-disk disruption in cardiac muscle (Dash et al., 2017). We therefore analyzed z-disks in septin 2-depleted cardiomyocytes to determine if septin 2 is similarly required for *Drosophila* cardiomyocyte z-disc integrity. As shown in Figure 5A, z-discs labeled with an anti-α-actinin antibody in *septin2^2^* homozygous mutant cardiomyocytes were nearly identical to those from *w^1118^* controls, with no differences in α-actinin labeling intensity or localization pattern. We did note, however, an approximately 25% increase in spacing between z-disks indicative of longer sarcomeres in *septin2^2^* compared to *w^1118^* cardiomyocytes (Figure 5A,B). This increase in sarcomere length may underlie the overall dilation of septin 2 as well as Stim depleted hearts, and is therefore consistent with SOCE suppression resulting from loss of septin 2.

**Figure 5.**
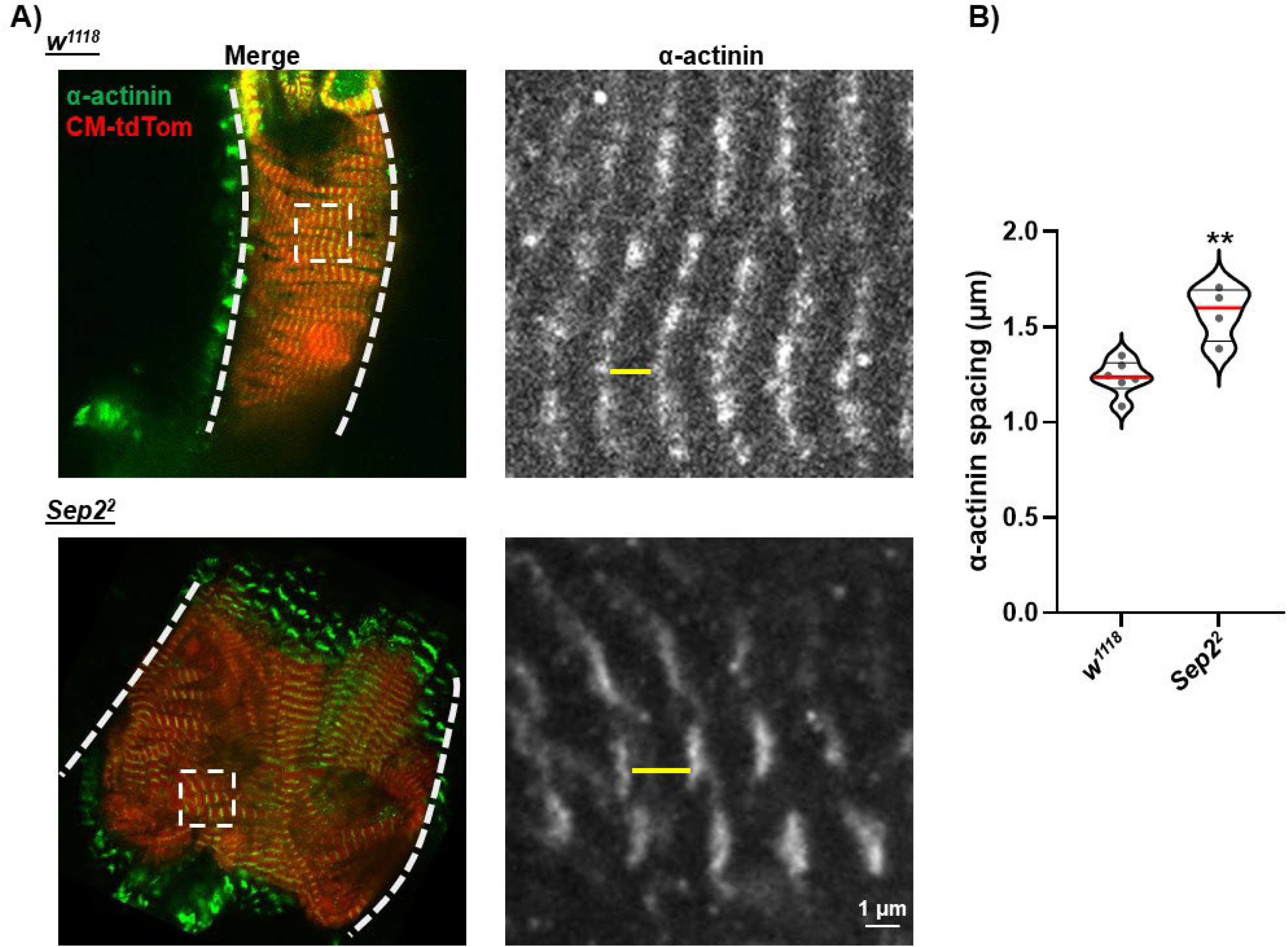
Septin 2 suppression results in increased z-disk spacing but does not alter z-disk integrity. A) Left: representative images of segments of hearts from *w^1118^* (control) or *Sep2^2^*animals labeled with anti-α-actinin antibody (green). Also shown is CM-tdTom fluorescence (red) to label cardiomyocytes. Images are z-projections encompassing approximately the dorsal half of the heart. Thick dashed lines indicate the lateral boundaries of the heart and the dashed boxed indicate the magnified regions shown on the right. Right: Magnified images of labeled α-actinin from boxed regions indicated in merged images on the left. Yellow lines indicate the spacing between adjacent z-disks. B) Plot of measured distance between α-actinin labeled z-disks from *w^1118^* control or *Sep2^2^* hearts. Each symbol represents the average of measurements from a single heart. Red lines indicate median and black lines indicate quartiles. **, P<0.01 compared to *w^1118^* control, two-tailed t-test.

## DISCUSSION

Collectively, our data demonstrate for the first time that cardiomyocyte expression of group 2 and group 6 septin monomers is required for proper heart function. We further show that the effects of septin suppression on heart function are likely due, at least in part, to suppression of cardiomyocyte SOCE. Cardiomyocyte-specific suppression of *Drosophila* group 2 septins 1 and 4 and group 6 septin 2 resulted in marked heart dilation and reduced fractional shortening, nearly identical to the effects of cardiomyocyte Stim and Orai suppression that we previously showed (Petersen et al., 2020). Direct genetic evidence of the functional association of septins with the SOCE machinery is provided by our data demonstrating that the effects of septin depletion are suppressed by SOCE upregulation through Stim^CA^ or Orai^CA^ expression. We also show that SERCA overexpression suppresses dilation associated with both STIM and Septin 2 depletion, further suggesting that both STIM and Septin 2 functions converge on the regulation of SR Ca^2+^ stores. Lastly we show that heart dilation due to Septin 2 depletion is not due solely to possible roles for septin 2 in heart development, and that septin 2 depletion does not result in disruption of α-actinin localization or organization at z-disks that anchor actin within thin filaments. While these latter experiments do not rule out all possible SOCE-independent functions of septins in heart physiology, they do reinforce our overall findings that SOCE regulation is an essential function of septins in cardiomyocytes.

The mechanisms by which septins regulate SOCE function in cardiomyocytes or other cell types are still not entirely clear. Data from mammalian cells demonstrate that septins do not directly interact with Stim or Orai proteins, but instead may regulate the microenvironment at E/SR-plasma membrane junctions to facilitate Stim-Orai interactions, as well as the number and size of the junctions themselves (Katz et al., 2019). Septins directly interact with membrane phospholipids (Bridges and Gladfelter, 2015; Szuba et al., 2021), and septin-dependent reorganization of PIP_2_ is required for Stim-Orai association and optimal Orai channel gating (de Souza et al., 2021; Sharma et al., 2013). Septins and PIP_2_ also coordinately regulate actin remodeling within ER-plasma membrane junctions in human HEK293 cells, and disruption of this remodeling through depletion of human SEPT4, PIP_2_, or the actin regulatory factors CDC42 and ARP2 disrupted STIM1 clustering at junctions and SOCE activation (de Souza et al., 2021). We attempted to analyze SR-plasma membrane junctions as well as Stim clustering at these junctions in *Drosophila* cardiomyocytes within the intact heart, but the complex architecture of the heart made imaging these structures extremely challenging. Thus it remains to be determined whether the phenotypes we observed with septin depletion in *Drosophila* cardiomyocytes similarly reflect a requirement for septins in regulating the membrane architecture required for Stim-Orai association. An interesting extension of this is whether septins in cardiomyocytes also regulate the SR – t-tubule dyad junctions that are required for functional coupling between L-type Ca^2+^ channels and RyRs for EC coupling. We did not observe any severe deficits in overall contractility of the heart as would be expected if EC coupling were compromised, suggesting that dyad junctions remain intact in septin-depleted cardiomyocytes. One possibility is that Stim and Orai interact at SR-plasma membrane junctions in cardiomyocytes that are distinct from the dyad junctions that bring RyR and L-type Ca^2+^ channels in close proximity, as has been suggested in mammalian skeletal muscle (Michelucci et al., 2018).

Interestingly, while depletion of group 2 and 6 septins consistently results in SOCE suppression in multiple cell types from flies to mammals, septin 7 depletion results in gain of SOCE function in *Drosophila* neurons and human neural progenitor cells (Deb et al., 2020; Deb and Hasan, 2019; Deb et al., 2016). We did not test septin 7 in this current study, but these prior findings suggest that SOCE regulation by septins is subunit specific. One possible explanation for this subunit specificity is that it is related to the position of the septin subunits within the hexameric septin filament (Deb and Hasan, 2016). For example, it is possible that loss of group 2 or 6 subunits results in shortened but not fully disassembled filaments, and that these truncated filaments interfere with PIP_2_ and actin regulation at membrane contact sites. Septin 7 depletion, on the other hand, may result in complete disassembly of filaments due to septin 7 occupying the central core positions within the filament. A complete lack of septin filaments may then allow dysregulated formation of ectopic membrane contact sites. Another possibility is that septin subunits may have functions that are independent of their assembly into filaments. This possibility is supported by the demonstration that septin 7 dimerization, which is required for filament assembly, is dispensable for septin 7 function in cytokinesis (Abbey et al., 2016).

Our evidence in support of septin regulation of SOCE in *Drosophila* cardiomyocytes would be further reinforced by direct measurements of SOCE using Ca^2+^ imaging in this system. Unfortunately, multiple attempts at these measurements in *Drosophila* cardiomyocytes from both intact and dissociated hearts were unsuccessful. However, direct SOCE measurements in isolated *Drosophila* neurons demonstrated that depletion of septin 2 or combined depletion of septins 1 and 4 resulted in near-complete suppression of SOCE-mediated Ca^2+^ influx (Deb and Hasan, 2019). Thus, septin regulation of SOCE is clearly conserved between *Drosophila* and mammalian cells. Direct SOCE measurements in isolated mammalian cardiomyocytes may offer further insight into the specific effects of septin depletion on cardiomyocyte Ca^2+^ transport and homeostasis.

In conclusion our results clearly demonstrate an essential role for septins in cardiomyocyte physiology and overall heart function. Our genetic data strongly support a specific role for septins in regulating the SOCE Ca^2+^ transport pathway. These findings further add to our understanding of the essential role that SOCE plays in regulating heart physiology, and suggest a novel functional association between cytoskeletal regulation and Ca^2+^ transport mechanisms in cardiomyocytes. Important future directions of this work include determining if septin regulation of SOCE is conserved in mammalian cardiomyocytes and directly analyzing the effects of septin disruption on cardiomyocyte Ca^2+^ transport and homeostasis.

## MATERIALS AND METHODS

### *Drosophila* Stocks and Husbandry

The following fly stocks were obtained from the Bloomington *Drosophila* Stock Center: *GAL4* control RNAi (35783), *STIM* RNAi (27263) *Septin 1* RNAi (27709) *Septin 2* RNAi (28004), *Septin 4* RNAi (31119), *SEP2^2^/TM6* (91003), *UAS-SERCA* (63228), and *TubP-GAL80[ts]* (7019). The *tinC-GAL4* stock was provided by Dr. Manfred Frausch (Freidrich Alexander University), and the CM-tdTom flies were obtained from Dr. Rolf Bodmer (Sanford Burnham Prebys Institute). *UAS-Stim^CA^*was generated by our lab as previously described (Petersen et al., 2022) and *UAS-Orai^CA^* was obtained from Dr. Gaiti Hassan (National Center for Biological Sciences, Bangalore, India). Flies were maintained on standard cornmeal fly food and crosses were at 25° C unless otherwise indicated.

### Heart dissection, Immunofluorescence labeling, and confocal imaging

Adult female flies were anesthetized with CO_2_ and adhered to a 20 mm petri dish dorsal side down with petroleum jelly. The flies were then bathed in an artificial *Drosophila* hemolymph solution (ADH) (108 mM NaCl, 5 mM KCl, 2mM CaCl2, 8mM MgCl2, 1mM NaH2PO4, 4 mM NaHCO3, 10mM sucrose, 5mM trehalose, and 5mM HEPES pH 7.1). The dissection proceeded by removing the head, the ventral section of the abdomen, and internal organs that obstructed the view of the heart. Once the heart was exposed, the ADH solution was replaced with new ADH containing 10 mM EGTA to stop heart contractions. Once the heart no longer exhibited visible contractions, the ADH with EGTA was removed and hearts were fixed with 4% paraformaldehyde (PFA) in PBS for 15 minutes. Fixed hearts were then washed three times for 10 minutes each with gentle rotation in PBS containing 0.1% Triton X-100 (PBS-T). The samples were then moved to a single well of a 96 well plate and incubated with primary antibody overnight at 4° C. The primary antibody used was mouse monoclonal anti-α-actinin (Developmental Studies Hybridoma Bank 2G3-3D7) at 1:1000. Samples were then washed three times in PBS-T for 10 minutes, and incubated in Alexa fluor 488 goat anti-mouse secondary at 1:500 for 1 hour at room temperature. After three 10 minute washes in PBS-T samples were mounted onto a glass microscope slide in Vectashield (Vectashield laboratories), with the dorsal exoskeleton facing the glass slide and the heart facing the coverslip. The samples were then imaged on a Nikon A1R confocal microscope using a 40X, 1.3 NA objective. Image stacks were acquired with a z-spacing of 1 µm to encompass approximately the dorsal half of the heart, and image stacks are presented as maximum intensity z-projections to show α-actinin labeling within single cardiomyocytes. Images were processed using ImageJ. α-actinin-labeled z-disk spacing was calculated by measuring the distance between adjacent z-disks using ImageJ.

### Intravital contractility imaging

Intravital imaging was conducted as previously described (Petersen et al., 2020). Briefly 3-7 day old female animals expressing CM-tdTom were anesthetized with CO_2_ and their dorsal abdomens were adhered to glass coverslips using Norland Optical Adhesive cured with a 48-watt UV LED light source for 60 seconds. Animals were allowed to recover for several minutes prior to imaging until they exhibited visible leg movements. CM-tdTom-labeled hearts were imaged through the dorsal cuticle on a Nikon Ti2 inverted microscope controlled with Nikon Elements software. Images were acquired at a rate of 200 frames per second using an ORCA-Flash4.0 V3 sCMOS camera (Hamamatsu) and excitation of CM-tdTom by 550 nm light from a Spectra-X illuminator (Lumencor). M-mode traces were generated using ImageJ by drawing a 1-pixel wide line through the A-1 chamber of the heart, and fluorescence intensity along this line was plotted over time using the MultiKymograph function of ImageJ. EDD and ESD were manually measured directly from the M-mode traces at points of full relaxation or contraction, respectively. The average of 5 measurements for EDD and ESD was reported for each animal and FS was calculated as [(EDD-ESD)/EDD] x 100. Heart rate was calculated from the M-mode traces by counting the number of contractions over the full timecourse.

### Conditional expression with TubP-GAL80[ts]

Fly crosses were carried out to obtain animals that carried *tinC-GAL4, TubP-GAL480[ts],* and UAS-RNAi transgenes. Crosses were maintained at 18° C until the time of eclosion of new progeny. Progeny with the appropriate genotypes were maintained at 18° C for an additional two days, and were then either kept at 18° C or transferred to 27° C for an additional 3-5 days. Animals were then analyzed by intravital heart imaging as previously described.

### Statistical analysis

Statistical analyses were carried out using GraphPad Prism software. Data sets with two conditions were analyzed with two-tailed t-tests and datasets with three or more conditions were analyzed by one way ANOVA with Tukey’s Multiple Comparisons. Statistical significance for all tests was considered at P < 0.05.

## ACKNOWLEDGEMENTS

This work was supported by Uniformed Services University intramural grant I80VP000404 and NIH grant R21 NS121821 to J.T.S. Reagents obtained from the Bloomington *Drosophila* Stock Center (NIH P40OD018537).

## DISCLAIMER

The opinions and assertions expressed herein are those of the author(s) and do not reflect the official policy or position of the Uniformed Services University of the Health Sciences or the Department of Defense.

