## Supplementary Data for "Septins regulate heart contractility through modulation of cardiomyocyte store-operated calcium entry"

### Supplementary Information

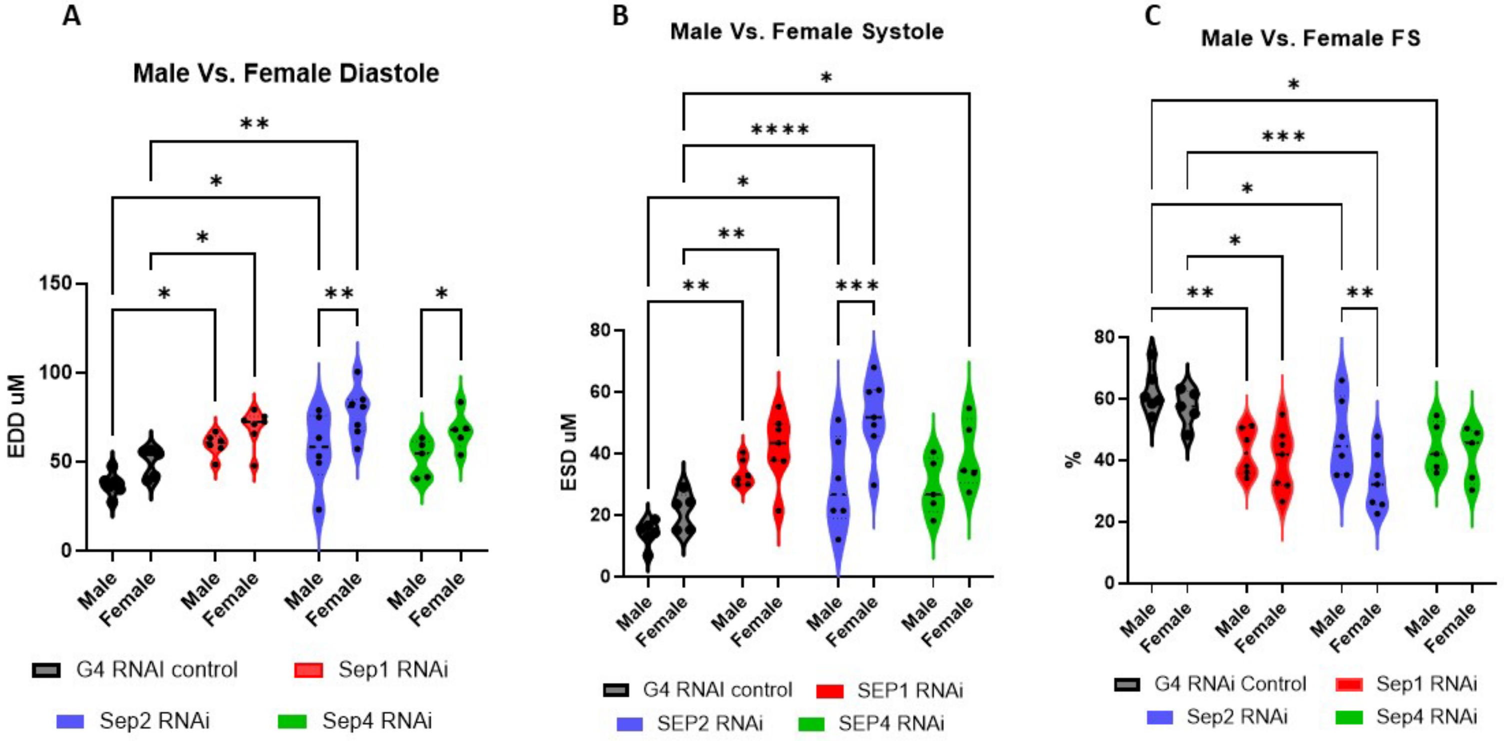


**<u>Supplementary Figure S1.</u> Septin depletion results in heart dilation in both male and female animals.** Shown are plots of EDD (A), ESD (B) and FS (C) for male and female animals with *tinC-GAL4* driven control or septin 1, 2, or 4 RNAi. Each symbol represents a single animal. *, P<0.05; **, P<0.01, ***, P < 0.001; ****, P < 0.0001, two-way ANOVA with Tukey's multiple comparisons.

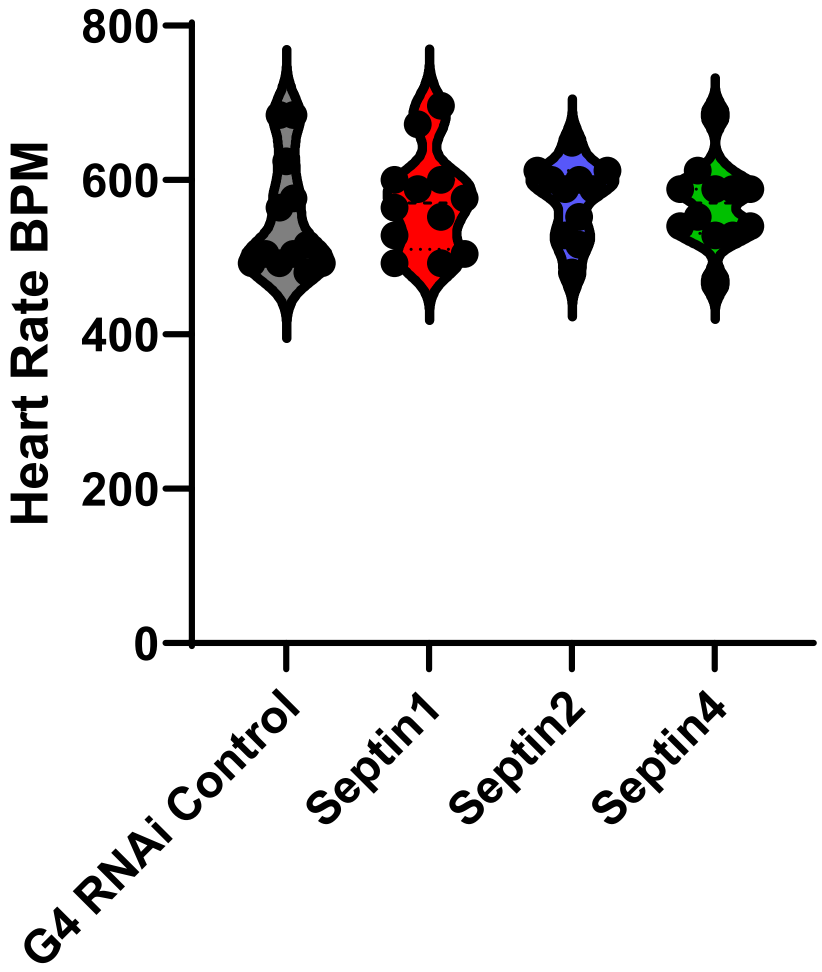


**<u>Supplementary Figure S2.</u> Septin depletion does not alter heart rate.** Shown are plots of heart rate in beats per minute (BPM) calculated from intravital imaging M-mode traces for animals with *tinC-GAL4* driven control or septin 1, 2, or 4 RNAi. There were no statistically significant differences; one-way ANOVA with Tukey's multiple comparisons.

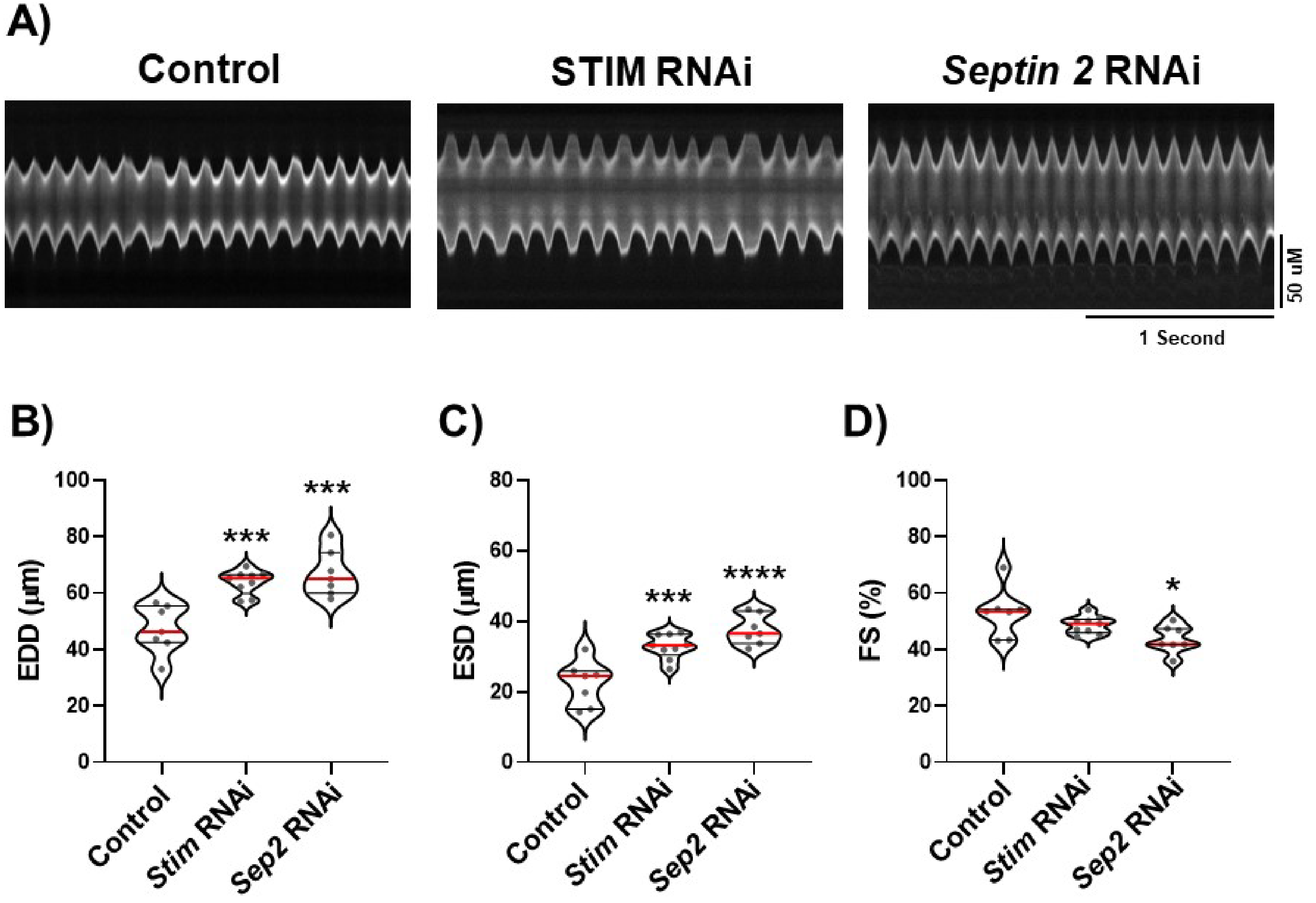


**<u>Supplementary Figure S3.</u>** ***Hand4.2-GAL4*** **driven** ***Stim*** **and** ***Septin 2*** **RNAi result in heart dilation.** A) Representative M-mode traces from animals with *Hand4.2-GAL4* driven control, *Stim*, or *Septin 2* RNAi. B) Plots of EDD, ESD, and FS, respectively, calculated from M-modes for animals with indicated *Hand4.2-GAL4* driven RNAi. Each symbol in plots represents the average of five measurements from a single animal's M-mode. Red lines indicate median and black lines indicate quartiles. *, P<0.05; **, P<0.01, ***, P < 0.001; ****, P < 0.0001; one-way ANOVA with Tukey's multiple comparisons.
